# Altered molecular signaling pathways in the hippocampus of rhesus monkeys following chronic alcohol use

**DOI:** 10.1101/2025.08.12.669930

**Authors:** Tanya Pareek, John M. Vergis, Xiaolu Zhang, Donna M. Platt, Kathleen A. Grant, Robert McCullumsmith, Barbara Gisabella, Sinead M. O’Donovan, Harry Pantazopoulos

**Affiliations:** Program in Neuroscience, University of Mississippi Medical Center, Jackson, MS 39216; Department of Neurosciences, University of Toledo, Toledo, OH 43606; Department of Microbiology and Immunology, Louisiana State University Health Sciences Center, Shreveport, LA 71105; Division of Neurobiology and Behavior Research, Department of Psychiatry and Human Behavior, University of Mississippi Medical Center, Jackson, MS 39216; Division of Neuroscience, Oregon National Primate Research Center, Oregon Health & Science University, Portland, OR 97239; Department of Behavioral Neuroscience, Oregon Health & Science University, Portland, OR 97239; Department of Psychiatry, University of Toledo, Toledo, OH 43606; Department of Biological Sciences, University of Limerick, Limerick, Ireland

## Abstract

Context-induced relapse is a significant factor limiting recovery from alcohol use disorder (AUD). However, the molecular processes in the hippocampus, a critical region for contextual memory impacted by chronic alcohol use, remain poorly understood. We used a non-human primate model to test the hypothesis that chronic alcohol use impacts hippocampal molecular pathways that may serve as therapeutic targets for context-induced relapse and memory processing issues associated with chronic alcohol use. We conducted RNAseq profiling on hippocampal samples from adult male rhesus monkeys with chronic alcohol use (n=7) and controls (n=5) from the Monkey Alcohol Tissue Research Resource (MATRR). We identified 2,575 differentially expressed genes (DEGs) in subjects with chronic alcohol use, including genes implicated in genome-wide association studies (GWAS) of alcohol dependence, such as GLP2R and GABBR2. Downregulated pathways included chemical synaptic transmission, trans-synaptic signaling, and neuron development, and upregulated pathways involved mitochondrial function. Targeted pathway analysis highlighted significant downregulation of synaptic signaling (e.g., axonal fasciculation) and upregulation of mitochondrial processes (e.g., electron transport). Leading-edge gene analysis revealed several downregulated genes involved in synaptic signaling including GRIN2B, CACNA1C, and NLGN1 as well as upregulated genes such as NDUFS3 and MT-ND1 involved in mitochondrial processes. Drug repurposing analysis identified several targets including epidermal growth factor receptor (EGFR) inhibitors, and L-type calcium channel blockers as potential therapeutic targets. Our results provide critical insights into molecular pathways underlying hippocampal pathology in chronic alcohol use, emphasizing the roles of mitochondrial function, synaptic regulation and calcium channels, and offering potential novel therapeutic targets.

## 1. INTRODUCTION

According to the 2023 National Survey of Drug Use and Health survey, approximately 28.9 million individuals in the United States were affected by an alcohol use disorder (AUD). Although available pharmacotherapies for AUD exist, high rates of relapse persist due to various complex factors [1]. Moreover, these treatments are effective only for certain populations [2], which highlights a need for the development of effective therapeutic strategies and novel pharmacotherapies to reduce relapse. There is a critical need to identify the molecular signaling pathways involved in chronic alcohol use and relapse in order to identify therapeutic targets for novel pharmacotherapies.

Drug relapse can be induced by a multitude of factors, including contextual cues and stress [3,4], which contribute to the formation and strengthening of drug-associated contextual memories. Furthermore, subjects with chronic alcohol use display enhanced memory for contexts associated with alcohol [5]. The hippocampus is at the center of neurocircuitry involved in reward memory processing [6], including contextual information associated with reward memories [7–9]. Specifically, accumulating evidence suggests that hippocampal subfield CA1 integrates dopamine reward signaling with contextual memories and regulates reward expectation and extinction [10,11]. For example, exposure to alcohol can enhance future alcohol intake by altering contextual and cue hippocampal processes [12], highlighting the importance of acute changes (i.e., enhancement of cue, context and reward processes) underlying learning processes following alcohol exposure. However, long-term exposure to alcohol can impair learning and memory by disrupting the hippocampal processes [13,14], and may contribute to generalized reward cues promoting relapse [15–18]. While acute and chronic alcohol exposure may cause differential effects in the hippocampus, such as enhanced contextual alcohol reward memory compared with general memory impairments, it is clear that hippocampal circuits play a critical role in relapse. Therefore, improving our understanding of molecular factors in the hippocampus involved in chronic alcohol use may provide potential targets for the development of effective therapeutic approaches for AUD.

Although pharmacotherapies are available for alcohol use disorder (AUD), effective treatments for context-induced relapse have not been identified. Furthermore, there is a lack of information regarding hippocampal alterations in chronic alcohol use in the primate hippocampus, limiting the development of novel therapeutic strategies. Current information from a small number of human studies is complicated by confounding factors inherent in human postmortem cohorts, such as comorbities and exposure to medications [19,20]. We used a nonhuman primate model to examine the effects of chronic alcohol use on molecular pathways of the primate hippocampus without the typical confounding factors present in human postmortem studies **(Figure 1)**. Our bioinformatic approach also identified potential therapeutic targets that may reverse the gene expression signatures identified in subjects with chronic alcohol use.

**Figure 1:**
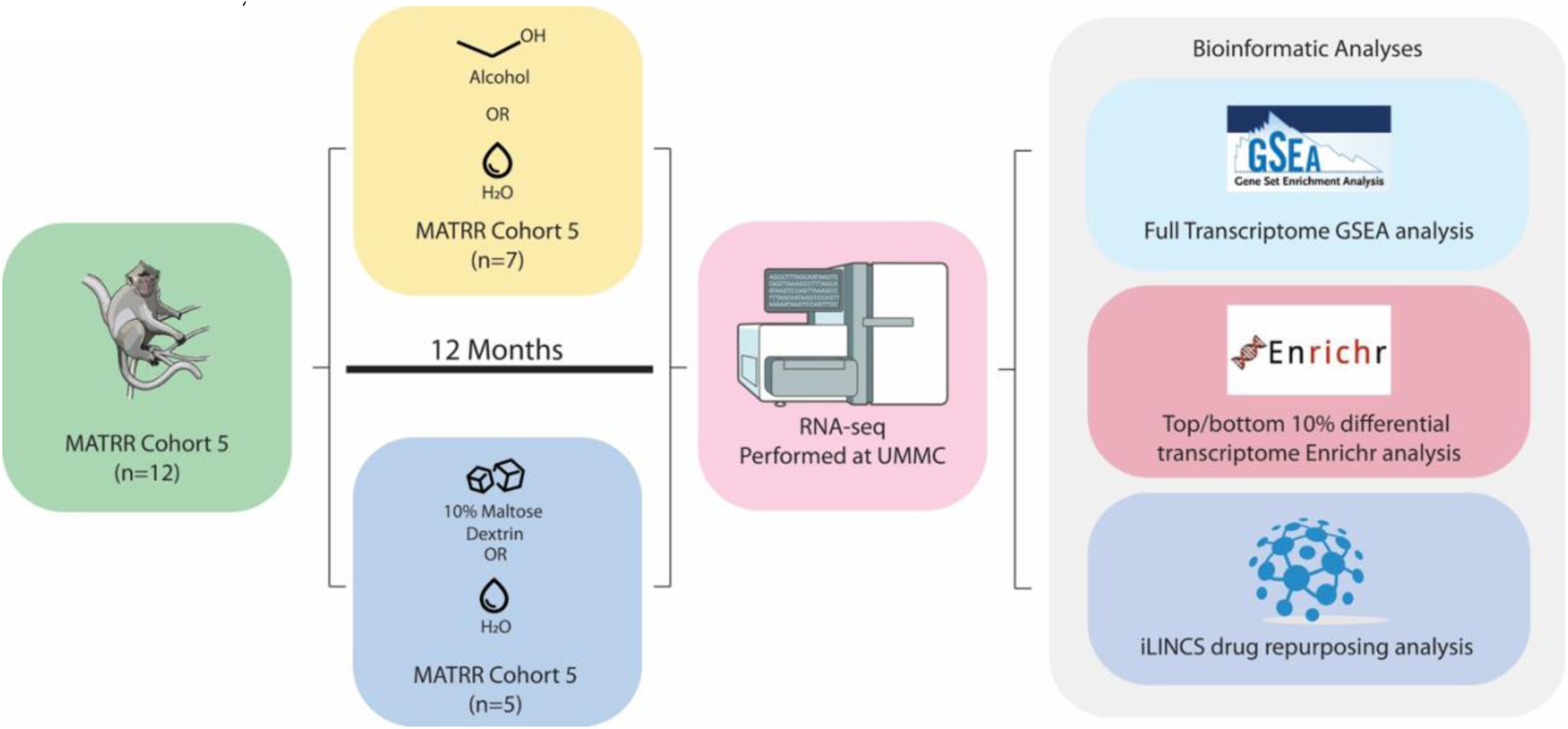
Study Overview. Twelve male rhesus monkeys from MATRR cohort 5 were given an open-access period with either ethanol and water (n=7), or 10% maltose dextrin or water (n=5) for 12 months. RNA-seq processing was performed at the UMMC Molecular Genomics Core. Furthermore, bioinformatic analyses were performed including GSEA pathway analysis, EnrichR pathway analysis, and iLINCS perturbagen analysis.

## 2. MATERIALS AND METHODS

### 2.1 Subjects and Tissue Collection

Hippocampal samples containing mid-body hippocampus were obtained from the Monkey Alcohol Tissue Research Resource (MATRR) Cohort 5 (www.matrr.com), see **Table S1** for demographic and experimental information. Fresh, frozen hippocampal samples from adult, male rhesus monkeys (*Macaca mulatta*) with a history of chronic, oral alcohol use (n=7; age= 7 yo) or no alcohol (n=5; age = 5-8 yo) were collected following the 12-month daily access to alcohol and water without imposed abstinence (i.e., within 2-4 hrs of the prior 22 hr self-administration session) [21]. All animal-use procedures were approved by the Oregon National Primate Research Center’s Institutional Animal Care and Use Committee and were conducted in accordance with the National Research Council’s Guide for Care and Use of Laboratory Animals (8th edition, 2011). Monkeys were experimentally naïve at the onset of alcohol induction and followed a standard operating procedure of alcohol self-administration [22]. Briefly, open-access (i.e., 22 hours/day) self-administration sessions occurred daily for 12 months, with average ethanol intake ranging between 2.55 and 3.78 g/kg/day. Cohort 5 monkeys used were defined as binge drinkers (BD; n=1), heavy drinkers (HD; n=2) or very heavy drinkers (VHD; n=4). Control monkeys’ self-administration sessions were identical to the alcohol oral self-administration sessions. However, the control monkeys had access to water and 10% maltose dextrin solution.

### 2.2 RNA sequencing

Hippocampal samples were removed from slides with sterile scalpels and processed for RNAseq profiling. RNA isolation, library preparation, and next-generation sequencing were performed by the Molecular and Genomics Core Facility at the University of Mississippi Medical Center, as described previously [23]. Total RNA was extracted from tissue samples using the Invitrogen PureLink RNA Mini kit with Trizol (Life Technologies; Carlsbad, CA, USA) according to the manufacture’s protocol. The quality control of the total RNA was evaluated with the Qiagen QIAxcel Advanced System and Qubit Fluorometer for quality and concentration measures, respectively.

Library preparation employed the TruSeq Stranded Total RNA LT Sample Prep Kit from Illumina (San Diego, CA, USA), per manufacturer’s protocol using up to 1 ug of RNA per sample. Libraries were index-tagged and pooled for multiplexing and sequenced on the Illumina NextSeq 500 platform with paired-end reads (2 x75 bp) using the Illumina 150 cycle High-Output reagent kit. Sequence reads were aligned to MacaM genome with the basespace application RNA-Seq Alignment (Version: 2.0.1 [workflow version 3.19.1.12+master]) that conducted both splice aware genome alignment with STAR alignment (version 2.6.1a, [24]) and transcriptome quantification with Salmon (version 0.11.2, [25]). RNAseq data are available at NCBI GEO GSE297855.

### 2.3 Differential gene expression analysis

Transcriptome-wide gene counts were subject to differential gene expression analysis between control and ethanol subjects using DEseq2 R package [26] with recommended default settings. Genes where there are less than 50% samples with normalized counts greater than or equal to 1 were filtered out.

### 2.4 Full transcriptome pathway analysis

#### 2.4.1. Gene Set Enrichment Analysis (GSEA)

with full set of genes was performed using fgsea R package (version 1.16.0) against human enrichment map gene sets downloaded from (https://baderlab.org/EM_Genesets/). sign(logFC) * (-log10(p-value)) or “stat” obtained from DESeq2 output were used as a gene ranking metric. Human genes were converted to rhesus monkey homologues as needed. GSEA first ranked the genes based on a measure of each gene’s differential expression, and then an enrichment score was generated (i.e., degree of overrepresentation of genes in a gene set at the top or bottom of the list of all ranked genes between ethanol and control conditions [27]. Unless otherwise specified, significantly altered genes or pathways (*p* <= .05) are represented by normalized enrichment score (NES). Here, a positive NES and negative NES represent a significant upregulation or downregulation, respectively.

#### 2.4.2 GSEA analysis also identified leading-edge (LE) genes using the leading-edge gene

analysis. Here, the focus was on genes that were represented most consistently across the top pathways within the GSEA analysis. Therefore, the subset of genes which can be interpreted as the core of a gene set that accounts for the enrichment signal [27]. Here, the overlap between multiple leading-edge subsets was analyzed.

### 2.5 EnrichR pathway analysis

GSEA with a gene set (i.e., top and bottom 10% genes; greatest absolute log2FC) was performed using enrichR R package (version 3.0). Gene ontology (GO) databases GO biological process, GO cellular component, and GO molecular function were used in analysis. Here, a combined score (CS) was generated to identify pathways with a high CS (i.e., a significant and robust enrichment unlikely due to randomness) or low CS (i.e., a weak enrichment may be due to randomness) that are significantly up or downregulated from a given list of DEGs [28].

### 2.6 iLINCS analysis

In order to identify perturbagens altering gene expression, the Library of Integrated Network-Based Cellular Signatures (LINCS) (https://www.ilincs.org/ilincs/) was used to identify concordant (i.e., causative) and discordant (i.e., treatment-related) gene signatures, mechanism of action (MOA), and pathways. LINCS is a National Institute of Health initiative that aims to create a comprehensive network of molecular reactions in response to environmental and internal stressors [29]. LINCS is a transcriptomic database that uses the L1000 assay, a gene expression array of 978 “hub” genes, to generate gene signatures. The L1000 genes are a reduced representation of the transcriptome, accounting for ∼82% of the information content of the transcriptome [30]. The database contains hundreds of thousands of gene signatures generated in human cell lines treated with chemical perturbagens (i.e., drugs) or following knockout or overexpression of individual genes. The log2 fold-change (log2FC) and p-value for the L1000 genes were extracted from DEG analysis and submitted as input to inquire a list of the chemical perturbagens altering the gene expression from the iLINCS portal. The reported score is the Pearson correlation coefficient between the disease signature and the precomputed iLINCS drug signatures. The chemical perturbagens with discordance scores < –0.321 and concordance scores > 0.321 were retained. Chemical perturbagens were clustered by MOA categories inquired from L1000 FWD [31], DrugBank database [32] and the Broad Institute.

### 2.7 Overlaps between GSEA and EnrichR pathways

Shared GO pathways between full transcriptome pathway analysis (GSEA) and targeted pathway analysis (EnrichR) were identified using the R script available at https://zenodo.org/badge/latestdoi/642681935.

## 3. RESULTS

### 3.1 Differential gene expression analysis

We identified 2,575 differentially expressed genes (DEGs) in the hippocampus of monkeys with chronic alcohol use with *p*-value < 0.05, with 553 DEGs meeting the criteria of *p*-adjusted significance (**Figure 2, Table S2**). These include several genes implicated in GWAS studies of alcohol dependence (e.g., GLP2R, GABBR2) [33]. Downregulated genes included GLP2R, GABBR2, genes involved in synaptic signaling (e.g., GRIN2A, GABRB2, KCNA1), genes involved in circadian rhythms and sleep (e.g., PER2, GSK3B, and ADORA1; **Figure S1, Table S2**) and extracellular matrix molecules (e.g., NCAN, TNR, SPOCK1, CTSB, ADAM22 and ADAM23; **Figure S1, Table S2**). Upregulated genes included genes involved in metabolism (e.g., NDUFA1, ND3, COX5B, ATP5E, ATP5L) ribosomal processes (e.g., RPL19, RPS29, RPL34, RPL21) and oxidative stress (e.g., NUDT1, ROMO1, NDUFB2, NDUFB1).

**Figure 2:**
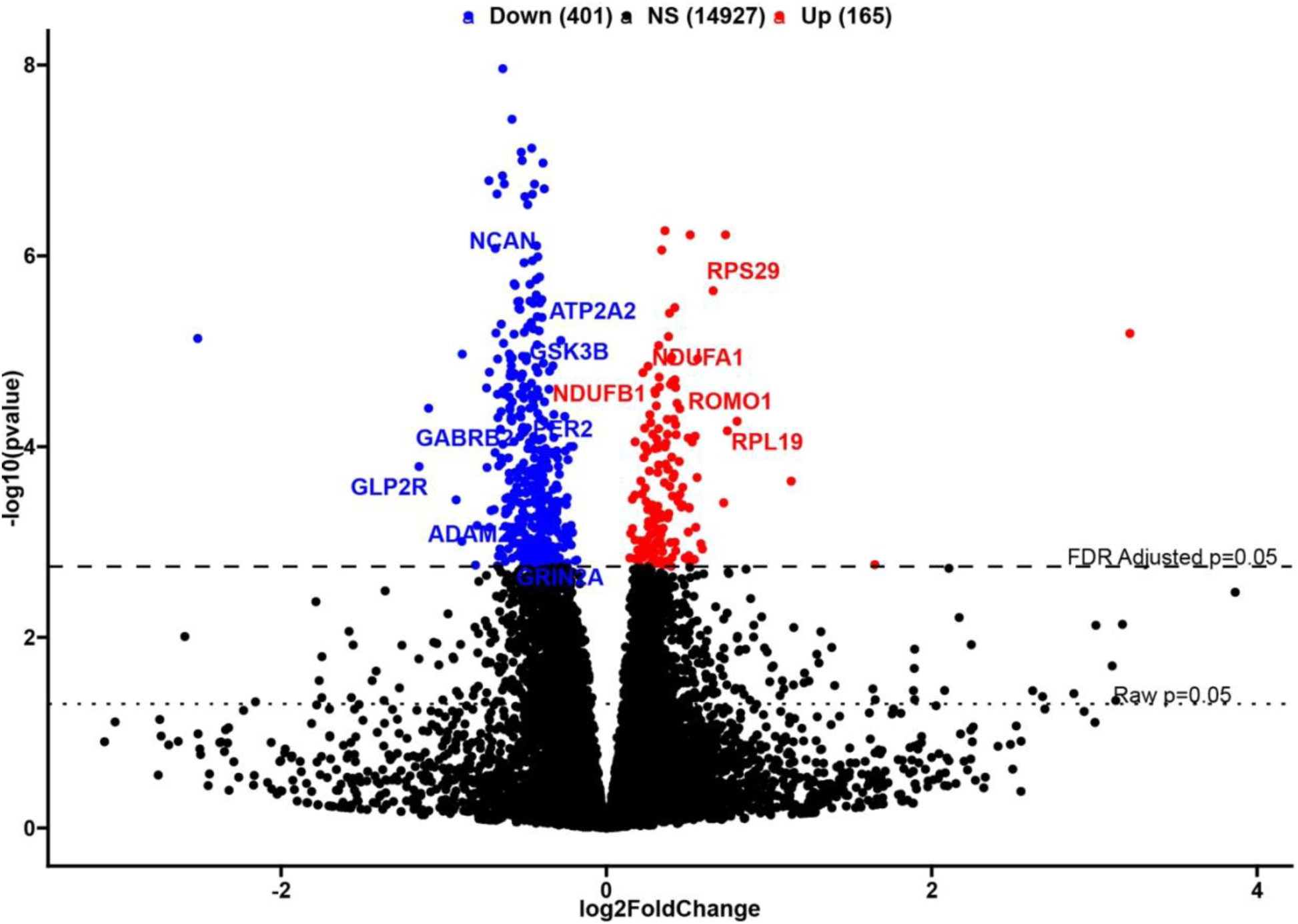
Volcano Plot of differentially expressed genes (DEG) in male monkeys with a history of chronic alcohol use. Differentially expressed genes in the hippocampus of monkeys with chronic alcohol use including several downregulated genes (e.g., GLP2R, GRIN2A) and upregulated genes (NDUFA1, RPS29). X-axis represents Log2 Fold Change (i.e., 1 or –1 signifies a 2-fold increase or decrease in gene expression, respectively). Y-axis represents –log10(p-value). Blue and red illustrate significantly (padj <0.05) down and upregulated genes, respectively.

### 3.2 Full transcriptome pathway analysis

#### 3.2.1 GSEA pathway analysis identified a total of 117

significantly altered pathways between the ethanol and control groups. Upregulated pathways, (57 total), included ribosomal process, metabolic processes and immune response (**Figure 3A, Table S3**). In comparison, downregulated (60 total) include pathways include regulation of trans-synaptic signaling, modulation of chemical synaptic transmission, synapse organization, postsynaptic density, dendritic spine, and regulation of synapse organization (**Figure 3A, Table S4**). The top 20 upregulated pathways include pathways related to cellular protein synthesis and energy metabolism (e.g., mitochondrial function; **Figure 3B**). Downregulated pathways include those involved in neuronal development, activity and synaptic signaling (**Figure 3B**).

**Figure 3:**
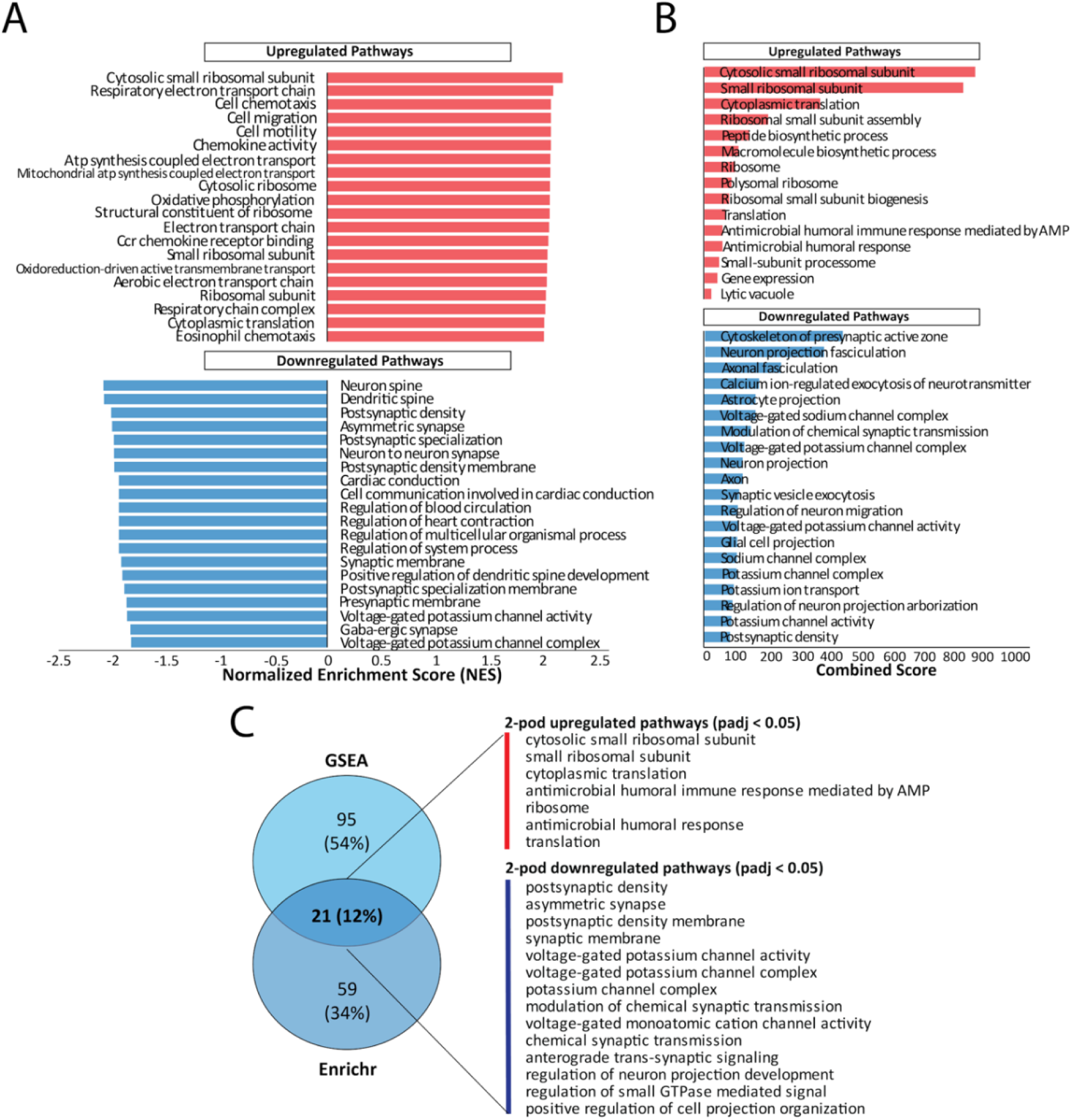
Pathway analysis. Full transcriptome pathway analysis of hippocampal pathways of monkeys with chronic alcohol use. **(A)** The top 20 upregulated pathways and downregulated pathways were plotted across normalized enrichment score on the X-axis. This score considers the overrepresentation of a gene set in a ranked list of all genes between two conditions (i.e., EtOH vs. control). A positive NES indicates significant upregulation at the top of the list, a negative NES signifies significant down-regulation at the bottom, while an NES close to 0 suggests no enrichment. Larger absolute values of NES imply stronger enrichment. **(B)** Targeted pathway analysis of top DEGs illustrated as by Enrichr including the top 20 upregulated pathways and downregulated pathways, where the X-axis represents the log combined score (i.e., logCS). The combined score (i.e., CS) ranks enriched pathways by multiplying the log of the enrichment analysis’s p-value (statistical significance) by the Z-score (degree of enrichment). A high CS denotes significant and robust enrichment unlikely due to random chance, whereas a low CS indicates weak enrichment or randomness. **(C)** A venn diagram of those pathways that are unique to either GSEA or EnrichR or those that are common between the two is depicted, along with a list of the 21 shared pathways.

### 3.3 EnrichR pathway analysis

EnrichR pathway analysis focuses on the top 10% and bottom 10% DEGs. The combined score (CS) values represent likelihood that enriched pathways are significant and unlikely due to chance. EnrichR analysis detected 81 significantly altered pathways with 15 upregulated pathways and 66 downregulated pathways, including calcium ion-regulated exocytosis of neurotransmitter and cytoskeleton of presynaptic active zone. Further, ribosomal pathways demonstrate higher expression levels in monkeys following chronic alcohol use, while synaptic signaling related pathways illustrate lower expression levels (**Figure 3B, Tables S7 and S8**), highlighting possible disruptions in communication between neurons. The top 20 upregulated pathways largely centered on the fundamental processes of protein synthesis, ribosome assembly, and antimicrobial humoral response (**Figure 3B**). Downregulated pathways consisted of pathways involved in neuronal structure, synaptic organization, and neurotransmission (e.g., calcium ion-regulated exocytosis of neurotransmitter, axonal fasciculation and postsynaptic density) (**Figure 3B**).

### 3.4 Overlaps between GSEA and EnrichR pathways

Overlap between the GSEA and EnrichR analyses, representing the most consistent changes identified across both analyses, identified 21 shared pathways. Shared upregulated pathways included small ribosomal unit, cytoplasmic translation, and antimicrobial humoral response (**Figure 3C**). Shared downregulated pathways included postsynaptic density, voltage gated potassium channel activity, and chemical synaptic transmission (**Figure 3C**).

### 3.5 Leading-edge gene analysis

Leading edge gene analysis identified transcripts that appear most frequently across the top and bottom pathways (i.e., their expression level increases or decreases in the chronic alcohol group compared to controls, respectively). The top 20 common leading-edge genes derived from our GSEA analysis (**Figure 4, Tables S5 and S6**) include statistically significant genes involved in inflammatory response (e.g., CCL2, CCL3, CCL11, C5 and IL6) and genes involved in mitochondrial function such as mitochondrial respiration, ATP production, and oxidative phosphorylation (e.g., NDFUFA2, NDFUFA5, NDFUFA8, NDUFB7, NDUFS4, MT-CYB, COX4I1, COXB5). Leading-edge genes in downregulated pathways included genes involved in synaptic transmission (e.g., NGLN1, GRIN2A, GRIN2B, DLG4, SHANK3), L-type voltage-gated calcium channel signaling (e.g., CACNA1C, CACNA2D1, and CACNA1G) and extracellular matrix molecule (NGLN1, NRXN1, TNR, ADAM22) are downregulated following chronic alcohol use, highlighting potential dysregulation.

**Figure 4:**
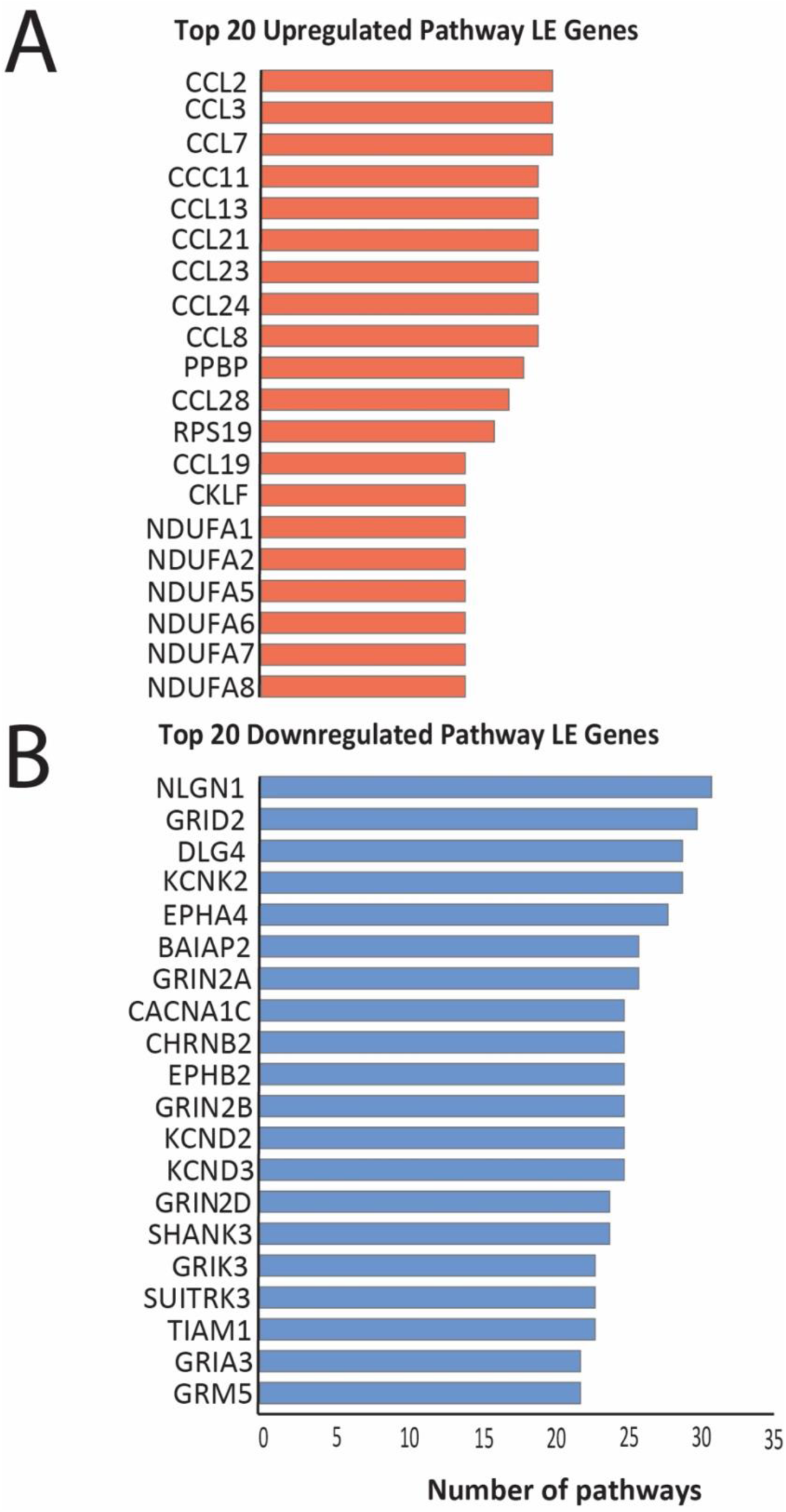
Leading Edge Gene Analysis. Top 20 leading edge genes **(A)** and bottom 20 leading edge genes **(B)** in the hippocampus of monkeys with chronic alcohol use. X-axis represents the number of pathways evaluated.

### 3.6 Drug Repurposing analysis using iLINCS

Signature-based connectivity analysis utilizing the Library of Integrated Network-based Signatures (LINCS) database was used to identify chemical perturbagens that reverse (discordant) or simulate (concordant) the transcriptional signatures we identified in the hippocampus of monkeys with chronic alcohol use (**Figure 5**). We identified 304 unique concordant (i.e., perturbagens that may induce disease-associated gene expression changes) and 373 discordant (putative therapeutic) mechanisms of action (MoAs). The top discordant mechanisms of action (MoAs) identified included VEGFR inhibitors, PDFGR inhibitors, NFkB inhibitors, serotonin receptor antagonists, dopamine receptor antagonists, calcium channel blockers and NADH dehydrogenase inhibitors (**Figure 5B, Table S12**). Along this line, we identified several discordant chemical perturbagen signatures representing potential therapeutic compounds, including AKT Inhibitor III, GSK-3 Inhibitor II, the serotonin/norepinephrine transporter inhibitor and dopamine antagonist Amoxapine, the L-type Ca channel antagonist Felodipine, and several P13K inhibitors (**Figure 5A, Table S10**). Further, top concordant pathways included GABA-gated chloride ion channel activity, GABA-A receptor activity, and benzodiazepine receptor activity as expected, as well as 3’,5’-cyclic-AMP phosphodiesterase activity, ionotropic glutamate receptor activity and high voltage-gated calcium channel activity (**Figure 5C, Table S15**). In comparison, the top discordant pathways included several pathways involved in ion channel and receptor-mediated signaling, including GABA-gated chloride channel activity, adenylate cyclase-inhibiting serotonin receptor signaling, and extracellular ligand-gated monoatomic ion channel activity, as well as pathways involved in energy metabolism and mitochondrial function, including mitochondrial electron transport, mitochondrial respiratory chain complex I and NADH dehydrogenase activity (**Figure 5D, Table S16**). In addition, 454 unique concordant and 497 unique discordant gene targets were identified (**Tables S13 and S14**).

**Figure 5:**
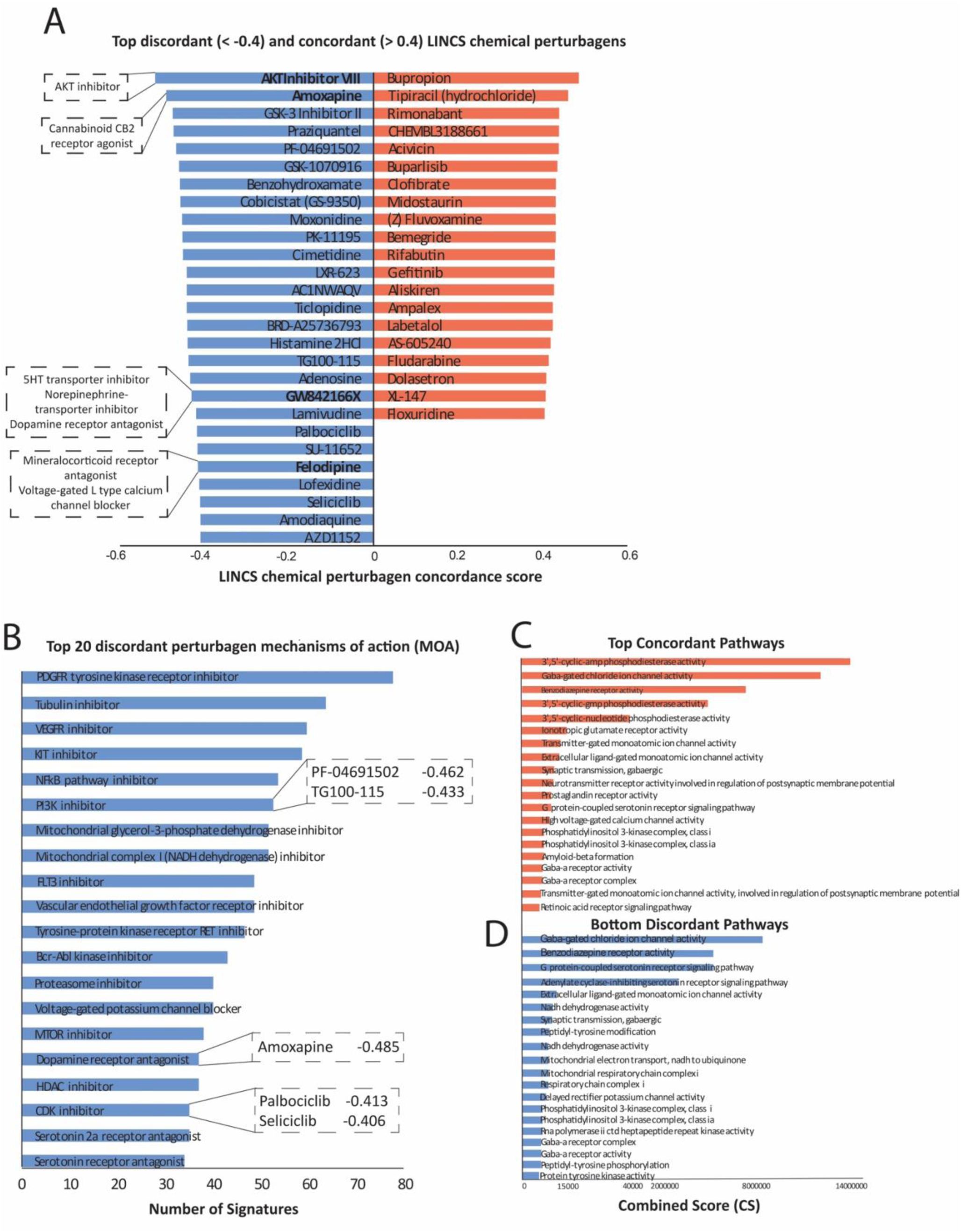
iLINCS Drug Repurposing Analysis. iLINCS drug repurposing analysis organized as top discordant (blue; < – 0.4) and concordant (red; > 0.4) chemical perturbagens **(A).** Here, X-axis represents the concordance (positive correlation coefficient) and discordance (negative correlation coefficient) scores. **(B)** illustrates the top 20 discordant mechanisms of action (MOA), where x-axis represents the number of signatures associated with a MOA. The 20 top and bottom pathways associated with concordant **(C)** and discordant **(D)** gene targets. CS represents the combined score in order to rank enriched pathways.

## 4. DISCUSSION

Although chronic alcohol use contributes to several hippocampal-dependent processes including memory impairment and context induced relapse, information regarding the molecular processes underlying contextual memory alterations in chronic alcohol use is currently limited, particularly in regard to humans and non-human primates. Furthermore, the information from a small number of human studies currently available is complicated by confounding factors inherent in human postmortem cohorts, such as comorbities and exposure to medications [19,20]. We used a nonhuman primate model to examine the effects of chronic alcohol use on molecular pathways of the primate hippocampus without the challenges of confounding factors present in human postmortem studies in order to identify effects specific to chronic alcohol exposure. Therefore, our dataset can also serve as a resource for interpreting potential confounding effects in current and future human postmortem studies. Our study represents to our knowledge the first molecular characterization of hippocampal alterations in non-human primates with chronic alcohol use. Dowregulated pathways involved in synaptic transmission and synaptic signaling along with upregulated pathways involved in mitochondrial processes and energy metabolism may contribute to memory alterations inherent in chronic alcohol use and AUD. Furthermore, our findings identify several potential therapeutic targets for context induced relapse.

Several synaptic signaling processes including chemical synaptic transmission, dendrite, presynapse, and regulation of synapse organization were amongst the most consistently identified downregulated pathways observed (**Figure 3A**), indicating broad synaptic hippocampal deficits following chronic alcohol use. Downregulation of synaptic signaling pathways identified by EnrichR pathway analysis, including cytoskeleton of presynaptic active zone, dendrite morphogenesis and GABA-ergic synapse pathways provides further support for broad synaptic deficits **(Figure 3B).** Furthermore, downregulated leading edge genes included genes in synaptic regulation such as NLGN1 and GRIN2A, GRIN2B, and GRIN1 **(Figure 4, Tables S5 and S6)**.

Downregulation of synaptic signaling pathways, including several glutamate receptors, indicates impaired hippocampal neurotransmission and synaptic plasticity. Glutamatergic NMDA receptors such as GRIN2A, GRIN2B and GRIN1 are essential for hippocampal learning [34,35]. Downregulated genes included several genes involved in synaptic regulation, including GLP2R and GABBR2 implicated GWAS studies of AUD [33]. Hippocampal decrease of GLP2R expression is associated with deficits in spatial memory processing, and upregulation of GLP2R enhances hippocampal neurogenesis and synaptic strength of hippocampal neurons [36], suggesting that the observed decrease of GLPR2 mRNA may contribute to hippocampal synaptic deficits with chronic alcohol use, and people with a GLP2R genetic predisposition may be more susceptible to such alterations. Broad decreases of synaptic signaling pathways may underlie memory deficits inherent with chronic alcohol use. Synaptic signaling decreases are in line with memory impairment reported in subjects with chronic alcohol use [16,18] as well as with preclinical studies reporting impaired spatial and contextual memory following alcohol exposure [37–39]. In addition to synaptic signaling, we detected downregulation of pathways related to chemical synaptic transmission, neuron projection development (e.g., axonogenesis and dendrite morphogenesis), and calcium ion transport, indicating a loss of neuronal connectivity. Such impairments in connectivity and plasticity have been implicated in the cognitive deficits observed in individuals with AUD [40–42].

Sleep and circadian rhythm disturbances are commonly reported in people with chronic alcohol use [43,44]. We observed the downregulation of genes involved in sleep and circadian rhythm regulation, including PER2, GSK3B and ADORA1 (see **Figure S1, Table S2).** Downregulation of the core clock gene PER2 is consistent with previous studies demonstrating that mice with a deletion in the PER2 gene display greater alcohol intake [45]. Furthermore, mutations in clock genes including PER2 have been associated with alcohol dependence [46]. Similarly, ADORA1 is critically involved in sleep regulation [47] with mice lacking ADORA1 expression displaying decreased sleep drive. Decreased hippocampal ADORA1 expression in monkeys with chronic alcohol use may contribute to insomnia associated with chronic alcohol use [48].

Our previous study using hippocampal samples from the same rhesus monkey cohort used in this study identified increased densities of perineuronal nets (PNNs) in monkeys with chronic alcohol use [49]. PNNs are extracellular matrix structures that form primarily around subsets of inhibitory interneurons, stabilizing synapses, including synapses involved in reward learning, and protecting neurons from oxidative stress [50]. We observed several downregulated extracellular matrix genes in the hippocampus of monkeys with chronic alcohol use including the endogenous proteases ADAM11, ADAMTS8, ADAMTS19, MMP16, CTSB (see **Figure S1, Table S2**), suggesting decreased PNN degradation in monkeys with chronic alcohol use. Increased PNNs in the presence of downregulated synaptic signaling pathways observed in the same subjects may reflect broad synaptic signaling decreases in the presence of stabilized alcohol reward-associated synapses by PNNs, possibly contributing to enhanced memory for contexts associated with alcohol reported in subjects with AUD [5]. Furthermore, PNNs in monkeys with chronic alcohol use may protect neurons from oxidative stress [51], as suggested by upregulated pathways involved in mitochondrial processes (**Figure 3A**).

In comparison, GSEA analysis identified fewer upregulated pathways than downregulated pathways (280 upregulated vs. 623 downregulated pathways) and largely consisted of mitochondrial processes, including protein synthesis, energy metabolism, oxidative phosphorylation and cellular respiration (**Figure 3A, Tables S3 and S4**). The top 20 leading-edge genes all consisted of genes involved in mitochondrial and NADH metabolic processes. Furthermore, EnrichR analysis identified more selective metabolic processes as the top upregulated pathways, including mitochondrial electron transport, RNA metabolic processes and NADH dehydrogenase activity. Upregulated pathways involved in protein synthesis and mitochondrial function may represent enhanced metabolic processes and increased oxidative stress induced by alcohol use [52,53]. Enhanced metabolic stress may reflect neurotoxicity from oxidative stress and inflammation reported in the brain following chronic alcohol use [54–56].

Our iLINCS perturbagen analysis identified putative therapeutic signatures and pathways encompassing dopamine and serotonin receptor antagonists, VEGFR/KIT inhibitors, PDFGR inhibitors, mitochondrial complex I inhibitors, and L-type calcium channel blockers **(Figure 5)**. Several pharmacological compounds are predicted to partially reverse the alcohol-induced gene expression changes in the hippocampus, representing potential pharmacological targets. These included pharmacological compounds such as AKT Inhibitor VIII, Amoxapine, GW842166X, and Felodipine **(Figure 5B)**. AKT activation has been implicated in prior alcohol use studies demonstrating that AKT activation in the nucleus accumbens contributes to alcohol self administration and binge drinking, which are attenuated by AKT inhibition [57]. Also, AKT has been implicated as part of a signaling pathway with PI3K and GSK3beta contributing to withdrawal effects from chronic alcohol exposure, and that PI3K inhibition can alleviate these effects [58]. Several PI3K inhibitors were identified as discordant chemical perturbagens in our analysis, including PF-04691502 and TG100-115 **(Figure 5C)**. Along the same line, the potent GSK3beta inhibitor GSK-3 Inhibitor II was identified as a top discordant chemical perturbagen **(Figure 5B)**. In addition to targeting the PI3K-AKT-GSK3beta pathway, GSK3beta is a core molecular clock gene, which may contribute to the molecular circadian clock gene alterations we observed **(Figure S1)**. Further, Amoxapine, a serotonin/norepinephrine transporter inhibitor and dopamine receptor antagonist, may be repurposed to address dysregulated dopaminergic signaling reported with chronic alcohol use [59,60]. Dopamine signaling to the hippocampus is critically involved in reward processing and relapse [61]. Hippocampal activity increases during reward anticipation, together with increased connectivity with the ventral tegmental area, indicating that dopamine modulation of the hippocampus by the VTA enhances hippocampal encoding during reward learning [62]. Furthermore, the hippocampus is critically involved in encoding predictability and uncertainty, key factors involved in reward processing. [63–65].

GW842166X is a cannabinoid CB2 receptor agonist **(Figure 5B)**. Several preclinical studies report that the CB2 receptor is involved in regulating the rewarding effects of alcohol [66,67]. CB2 receptor knockout mice demonstrate increased alcohol preference and consumption, and genetic polymorphisms for the CB2 gene have been associated with AUD [66]. Furthermore, CB2 receptor knockout mice display increased proinflammatory makers in the hippocampus in response to alcohol exposure, suggesting that CB2 agonists may also reduce inflammatory effects of alcohol on this brain region [67].

Felodipine is a mineralcorticoid receptor antagonist and L-type calcium channel (LTCC) blocker **(Figure 5B)**. Preclinical studies report increased L-type calcium channel expression and amplitude of LTCC currents in the hippocampus in alcohol dependent rats [68]. Furthermore, cue-induced alcohol reinstatement in these animals was attenuated by central administration of the LTCC antagonist verapamil [68], providing support for the use of these blockers as potential therapeutic agents. Our observed high voltage gated calcium channel activity concordant pathway **(Table S15)** provides additional support for LTCC blockers as a potential therapeutic strategy. In addition, felodipine has been reported to attenuate neuroinflammation [69], suggesting it may counteract multiple effects of alcohol on the hippocampus. Overall, these findings highlight several targets for future investigations into pharmacological agents to alleviate hippocampal pathology associated with chronic alcohol use.

Finally, converging evidence between the full transcriptome GSEA and EnrichR analyses highlighted shared upregulated pathways related to small ribosomal unit, cytoplasmic translation, and antimicrobial humoral response (**Figures 3C and S3**) and shared downregulated pathways involved in postsynaptic density, voltage gated potassium channel activity, and chemical synaptic transmission (**Figure 3C and S3**). The top shared pathways between these pods provides further support for decreased synaptic signaling and upregulated gene expression pathways promoting neuroinflammatory response and cell survival in response to enhanced mitochondrial stress. Impaired synaptic signaling and metabolic stress may underlie memory impairment and enhanced memory for contexts associated with alcohol [5] reported in subjects with chronic alcohol use [16,18].

Although our study provides a number of gene expression pathways and potential pharmacotherapy targets, our analysis was limited to bulk RNAseq. Future studies examining cell-type specificity and protein measures would provide insight into the neurocircuitry underlying our findings. Furthermore, our iLINCS perturbagen analysis was limited to gene expression signatures for compounds and cell types currently available in this database. Therefore, future studies using expanded databases may identify additional and more selective therapeutic targets for follow-up behavioral and longitudinal studies.

In summary, we identified a number of gene expression pathways altered in the primate hippocampus by chronic alcohol, including impaired synaptic signaling and metabolic stress that may contribute to memory impairment and enhanced memory for contexts associated with chronic alcohol use [5,16,18] along with several potential therapeutic drug targets for this hippocampal pathology. Future studies evaluating cell type specificity and the effectiveness of potential therapeutic compounds (e.g., AKT Inhibitor II, Amoxapine, PI3K inhibitors, and Felodipine) may provide insight into therapeutic targets for hippocampal dysfunction in subjects with chronic alcohol use.

## AUTHOR CONTRIBUTIONS

T.P. contributed to data collection, J.M.V., X.Z., and S.O. analyzed the data, T.P., H.P., S.O., B.G, D.M.P., K.A.G. and R.M. contributed to data interpretation, T.P., J.M.V., S.O. and H.P. wrote the manuscript, B.G., D.M.P., K.A.G. and R.M. contributed to manuscript preparation, H.P., T.P., S.O. and B.G. designed the studies,

