## Supplemental Materials for "Altered molecular signaling pathways in the hippocampus of rhesus monkeys following chronic alcohol use"

**Table 1.**

**Table 1. MATRR cohort information (Cohort 5)**

| Sex | Age | Group | Drinking Days | Drinking Category |
|---|---|---|---|---|
| **M** | 7 | EtOH | 364 | BD |
| **M** | 7 | EtOH | 365 | HD |
| **M** | 7 | EtOH | 356 | HD |
| **M** | 7 | EtOH | 365 | VHD |
| **M** | 7 | EtOH | 364 | VHD |
| **M** | 7 | EtOH | 363 | VHD |
| **M** | 7 | EtOH | 361 | VHD |
| **M** | 8 | Control | 0 | N/A |
| **M** | 8 | Control | 0 | N/A |
| **M** | 6 | Control | 0 | N/A |
| **M** | 5 | Control | 0 | N/A |
| **M** | 5 | Control | 0 | N/A |

Binge Drinker (BD), Heavy Drinker (HD), Very Heavy Drinker (VHD), Ethanol (EtOH)

**Figure 1. Circadian rhythm and extracellular matrix differentially expressed genes in male monkeys with a history of chronic alcohol use**

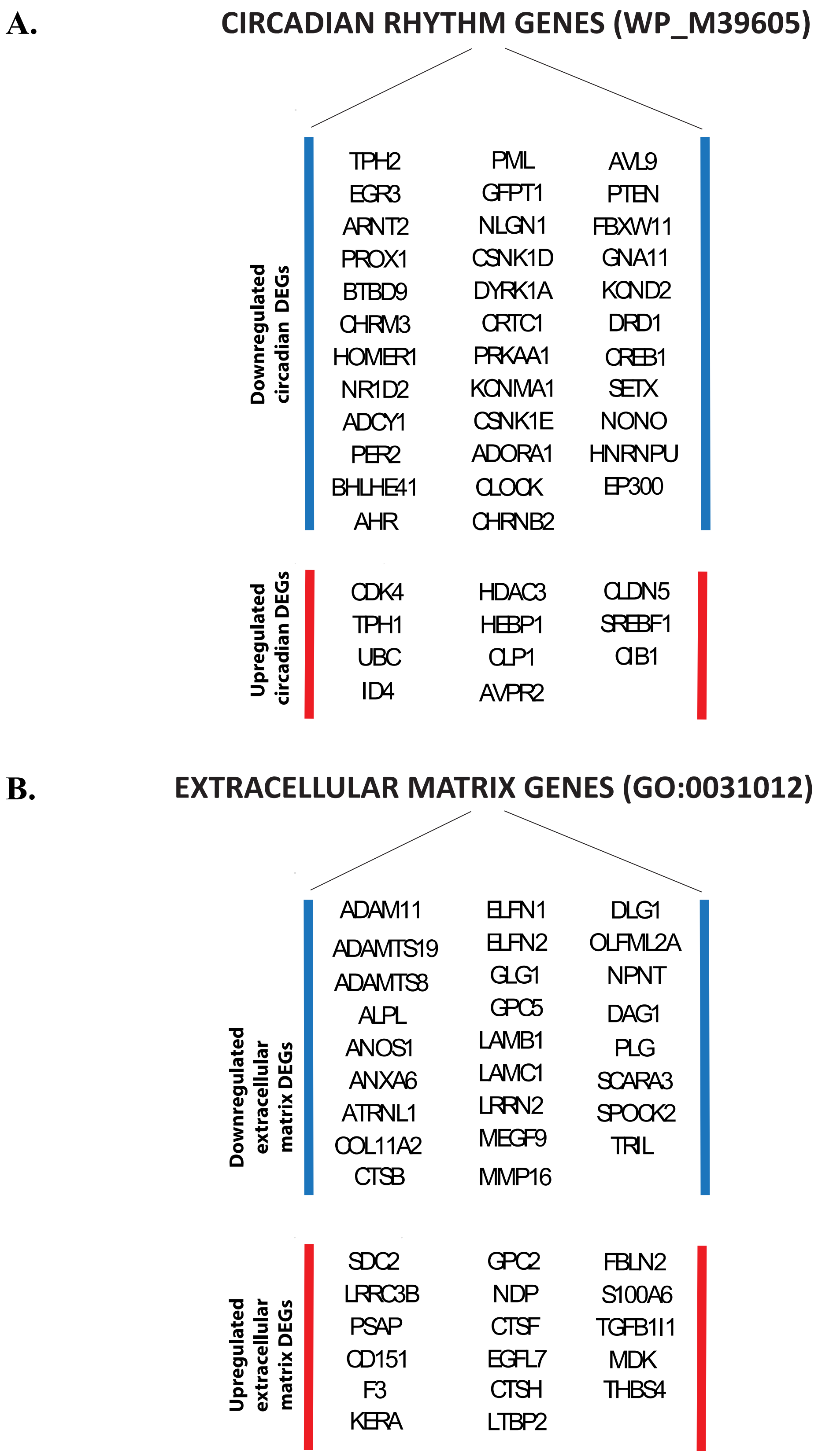

**Figure 2. LINCS analysis heatmap**

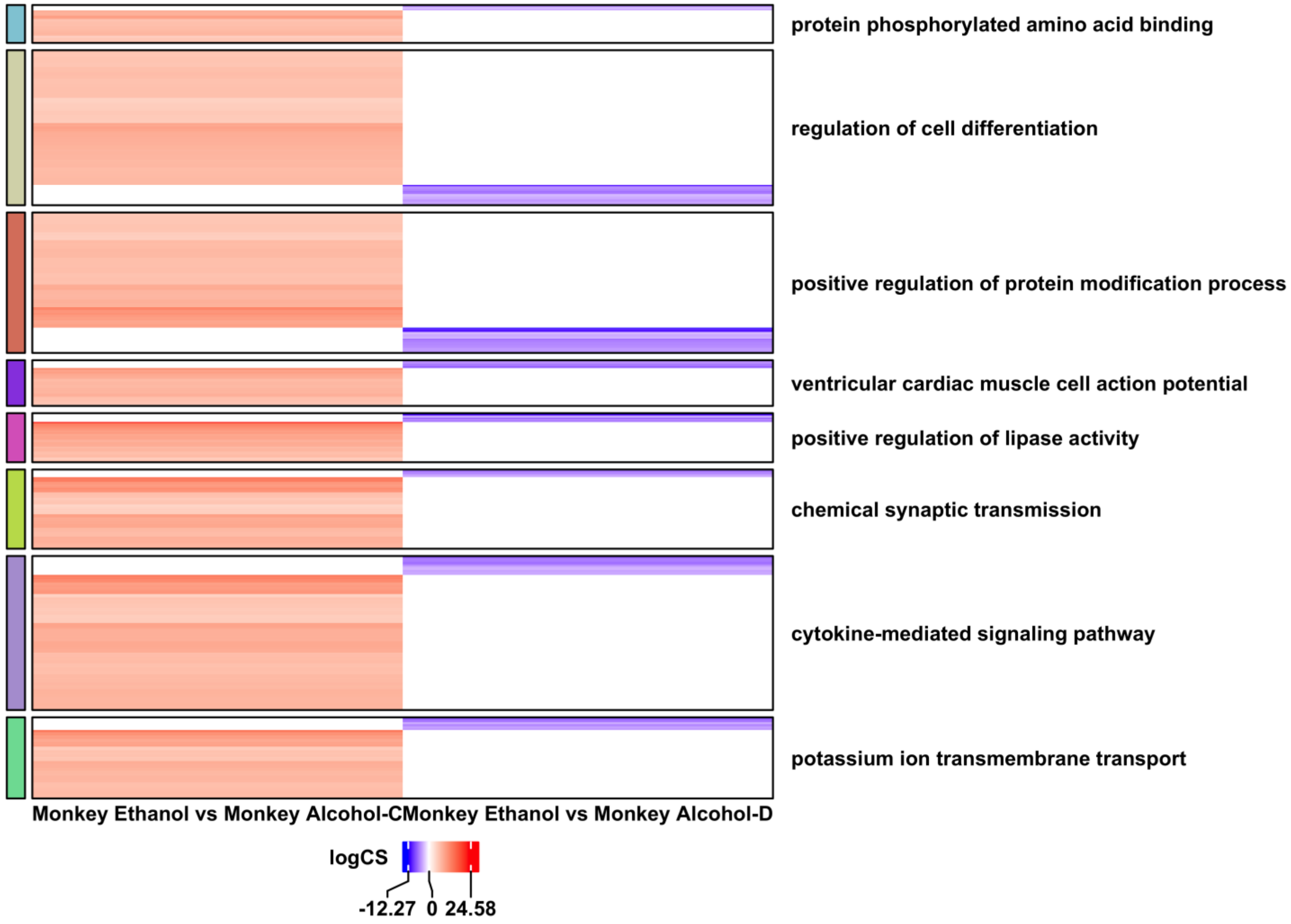

### Figure Captions

**Table 1. MATRR Cohort 5 demographic and experimental information**

**Figure 1. Circadian rhythm and extracellular matrix differentially expressed genes (DEG) in male monkeys with a history of chronic alcohol use** Blue and red on y-axis illustrate significantly down and upregulated genes, respectively.

**Figure 2. iLINCS perturbagen analysis** organized as a heatmap **(A);** x-axis represents the log combined score (i.e., logCS).

**Table 2. DEG analysis** All differentially expressed genes in the hippocampus of monkeys with alcohol use and no alcohol use.

**Tables 3-4. GSEA analysis** All upregulated and downregulated hippocampal pathways from full transcriptome pathway analysis.

**Tables 5-6. Leading edge gene analysis** All top and bottom leading edge genes in the hippocampus of monkeys.

**Tables 7-8. Enrichr analysis** All upregulated and downregulated pathways from focused analysis of top DEGs.

**Tables 9-16. iLINCS analysis** All concordant and discordant perturbagens, mechanism of actions, gene targets and pathways from perturbagen analysis.
